# Understanding the Ca^2+^-dependent Fluorescence Change in Red Genetically Encoded Ca^2+^ Indicators

**DOI:** 10.1101/435891

**Authors:** R.S. Molina, Y. Qian, J. Wu, Y. Shen, R.E. Campbell, T.E. Hughes, M. Drobizhev

## Abstract

Genetically encoded Ca^2+^ indicators (GECIs) are widely used to illuminate dynamic Ca^2+^ signaling activity in living cells and tissues. Various fluorescence colors of GECIs are available, including red. Red GECIs are promising because longer wavelengths of light scatter less in tissue, making it possible to image deeper. They are engineered from a circularly permuted red fluorescent protein fused to a Ca^2+^ sensing domain, calmodulin and a calmodulin-binding peptide. A conformational change in the sensing domain upon binding Ca^2+^ causes a change in the fluorescence intensity of the fluorescent protein. Three factors could contribute to this change in fluorescence: 1) a shift in the protonation state of the chromophore, 2) a change in fluorescence quantum yield, and 3) a change in the extinction coefficient for one-photon excitation or the two-photon cross section for two-photon excitation. We conducted a systematic study of the photophysical properties of a select cohort of red GECIs in their Ca^2+^-free and Ca^2+^-saturated states to determine which factors are most important for the Ca^2+^-dependent change in fluorescence. In total, we analyzed nine red GECIs, including jRGECO1a, K-GECO1, jRCaMP1a, R-GECO1, R-GECO1.2, CAR-GECO1, O-GECO1, REX-GECO1, and a new variant termed jREX-GECO1. We found that these red GECIs could be separated into three classes that each rely on a particular set of factors. Furthermore, in some cases the magnitude of the change in fluorescence was different depending on one-photon excitation or two-photon excitation by up to a factor of two.

## Introduction

The advent of fluorescent genetically encoded Ca^2+^ indicators (GECIs) has been revolutionary in understanding key aspects of biology, ranging from immunology (1, 2) to neuroscience (3, 4). These engineered proteins make it possible to visualize dynamic Ca^2+^ levels in genetically specific live cells and tissues under one-photon or two-photon excitation. There are many different GECI designs (5, 6). The most popular design is based on a single fluorescent protein (FP) that is circularly permuted in the middle of its 7th β-strand and fused to calmodulin (CaM) at the C-terminus and a calmodulin-binding peptide such as RS20 at the N-terminus. Initially, only green FP based sensors of this type were made (7–9), and their continuous development through several generations to increase Ca^2+^ sensitivity resulted in the widely used GCaMP6 series (10). Recently, the color palette was broadened to include blue and red (11). Red GECIs are especially desirable because red wavelengths of light scatter less in living tissue than green or blue, enabling deeper imaging (12). In addition, red-shifted excitation wavelengths are associated with less autofluorescence and photodamage.

The first red GECI, R-GECO1 (11), was based on the directed evolution of circularly permuted mApple (13) fused to the same Ca^2+^-sensing domain as GCaMPs. From the R-GECO1 template, protein engineering efforts through random mutagenesis as well as site-specific mutations inspired by various red FPs yielded the improved R-GECO1.2, a red-shifted variant CAR-GECO1, a long Stokes shift variant REX-GECO1, and a blue-shifted variant O-GECO1 (14, 15). The latter likely acquired the mOrange-type chromophore (16) from a Met to Thr mutation at the first amino acid in the chromophore-forming tripeptide (14). Other R-GECO1 based variants include RCaMP1.07 and RCaMP-2 (17, 18). One of the latest variants is jRGECO1a (19), which was produced through directed evolution with a neuron screening platform with the goal to better detect Ca^2+^-transients produced by neuronal action potentials. An entirely separate line of red GECIs based on a different circularly permuted red FP, mRuby (20), was created and designated the RCaMPs (21). RCaMPs have also been further improved with the neuron screening platform, resulting in jRCaMP1a and jRCaMP1b (19). Most recently, K-GECO1 was developed from a third circularly permuted FP, mKate-derived FusionRed (22, 23).

In this wide range of red GECIs, binding to Ca^2+^ causes a dramatic increase in fluorescence intensity. There are three factors that can contribute to a Ca^2+^-dependent fluorescence change on a molecular level. The first is a shift in the protonation state of the chromophore, as the protonated and deprotonated forms behave quite differently. The second is a change in the fluorescence quantum yield. The third is a change in the molecular absorption coefficient at the excitation wavelength, which is the extinction coefficient in the case of one-photon excitation or the two-photon cross section in the case of two-photon excitation. This last factor could depend on shifts of the excitation spectrum, and it could be different under one-photon and two-photon excitation.

In a detailed study of the popular green GECI GCaMP6m, Barnett et al. found that for this particular sensor the main mechanism of its Ca^2+^-dependent fluorescence increase was a shift in the protonation state of the chromophore (24). When it bound to Ca^2+^, the conformational change of the protein translated into a change in the electrostatic potential around the chromophore and a consequent shift in the chromophore p*K*_*a*_, pushing the equilibrium from the protonated, neutral form toward the deprotonated, anionic form. Since for GCaMP6m, only the anionic form fluoresces when excited at the wavelengths used to detect Ca^2+^, this resulted in an increase in fluorescence.

We conducted a similar systematic study for the range of red GECIs. Our goal was to investigate the photophysical properties of a cohort of available red GECIs to find the main factor(s) responsible for the Ca^2+^-dependent change in fluorescence under both one-photon and two-photon excitation.

## Background and Analytical Considerations

Figure 1A presents a typical red GECI design, using K-GECO1 in its Ca^2+^-saturated state (22) as an example. It consists of a red FP that is circularly permuted so that the original N-and C-termini are linked together and new ones are introduced into the 7th β-strand of the 11-β-strand barrel adjacent to the phenolic oxygen of the chromophore. The new termini are attached via short peptide linkers to a two-part Ca^2+^-sensing domain: the Ca^2+^-binding protein calmodulin (CaM) at the C-terminus and a calmodulin-binding peptide at the N-terminus. Depending on the red GECI, the N-terminus arm could either be the rat CaM-dependent kinase kinase peptide ckkap (as in K-GECO1 and R-CaMP2) or the chicken myosin light-chain kinase peptide RS20 (as in the other red GECIs mentioned). In a Ca^2+^-free state, the CaM and its binding peptide are apart, and the FP is dim. In a Ca^2+^-saturated state, CaM binds to four Ca^2+^-ions and undergoes a conformational change to wrap around the CaM-binding peptide, which results in a fluorescence increase. Since the entire construct is genetically encoded, it is possible to follow Ca^2+^-dynamics by expressing it in living systems such as neurons and imaging fluorescence over time (Figure 1B).

How can binding to Ca^2+^ modulate the fluorescence of the FP? As the chromophore is the molecule responsible for the fluorescence, we begin by examining its two possible protonation states, the protonated neutral form and the deprotonated anionic form (Figure 1C).

**Figure 1.**
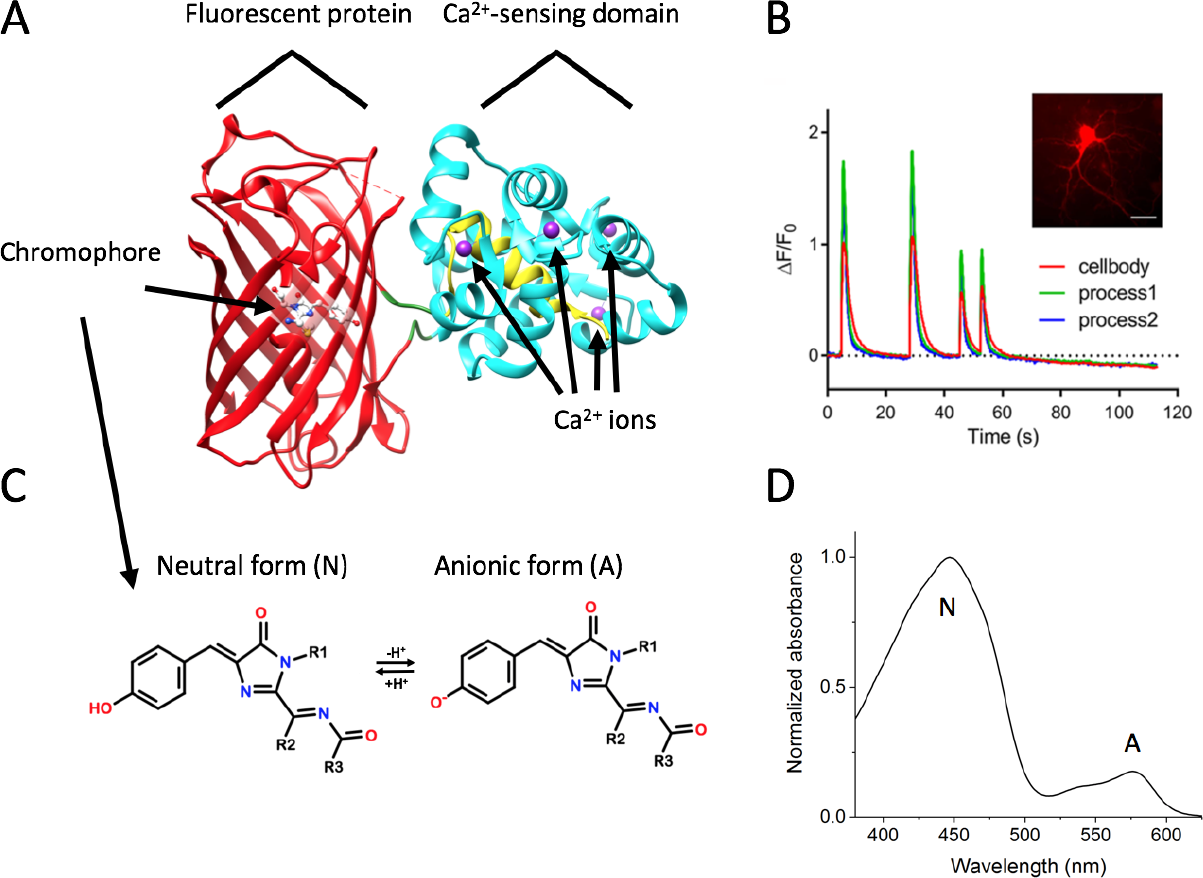
Red GECIs structure, function, and chromophore. (*A*) Crystal structure of K-GECO1 in the Ca^2+^-saturated state (PDB: 5UKG). The FP is colored red, CaM is cyan, the CaM-binding peptide (ckkap in K-GECO1) is yellow, and the Ca^2+^ ions are purple. (*B*) Imaging of spontaneous Ca^2+^ oscillations in dissociated rat hippocampal neurons with K-GECO1. Graph shows the change in fluorescence normalized to baseline fluorescence over time of three ROIs. Inset: Widefield image of K-GECO1 expression in neurons (scale bar, 30 μm). Adapted from (22). (*C*) Molecular structure of the typical red FP chromophore in its neutral and anionic forms. (*D*) Absorption spectrum of R-GECO1 in the Ca^2+^-free state illustrating the presence of both forms of the chromophore.

Nestled inside the FP β-barrel, the red FP chromophore exists in an equilibrium between its neutral and anionic forms that can shift in a Ca^2+^-dependent manner. The neutral (protonated) form maximally absorbs light at about 450 nm, while the anionic (deprotonated) form absorbs at about 570 nm (Figure 1D). Each form has a distinct fluorescence quantum yield (ϕ), extinction coefficient (ε), and two-photon cross section (σ_2_). These properties can change depending on whether the protein is bound to Ca^2+^ or not. Because of this, the neutral and anionic forms have their own one-photon brightness (ε×ϕ) and two-photon brightness (σ_2_×ϕ) which, weighted by their relative fractions (ϱ), combine to make the total brightness per molecule of a GECI in the Ca^2+^-saturated and Ca^2+^-free states. Equations 1 and 2 show the total one-photon brightness (F_1_) and two-photon brightness (F_2_) as a function of excitation wavelength (λ), where the subscripts A and N denote the anionic or neutral form, respectively, and the relative fraction ϱ_X_ is defined by the concentration of form X divided by the total concentration of chromophore.

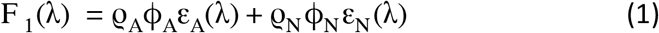

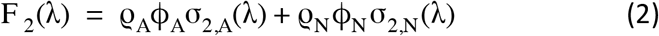

The fluorescence ratio between the Ca^2+^-saturated and Ca^2+^-free states can be expressed as F_n,sat_/F_n,free_, where the subscript n is 1 or 2. Conveniently, at the excitation wavelengths that are relevant to sensing Ca^2+^ (i.e. the F_n_ ratio is at or near maximum), only one form of the chromophore per Ca^2+^-state has to be considered because the brightness of the other form is close to zero. Here, the fluorescence ratios simplify to Equations 3 and 4, where the subscript e denotes the “excitable” form of the chromophore. The excitable form could be neutral or anionic, and it could also be different depending on the Ca^2+^ state.

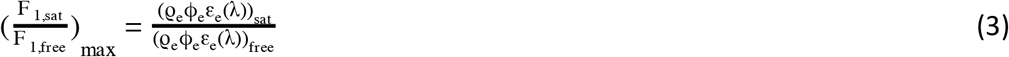

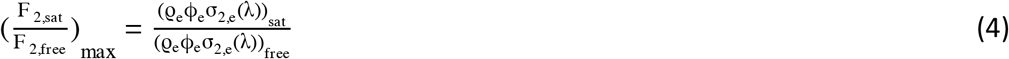

All three factors of the excitable form, (1) the relative fraction ϱ_e_, (2) the fluorescence quantum yield ϕ_e_, and (3) the molecular absorption coefficient ε_e_ or σ_2,e_, can change when the protein binds to Ca^2+^. With careful measurements of both the Ca^2+^-saturated and Ca^2+^-free states of each GECI, we can quantify each factor to discover which one(s) are the most important.

Because there are two forms of the chromophore, measuring the ϕ corresponding to just one form can require a more involved approach. Specifically, when the excitable form of the chromophore is anionic but the concentration of the neutral form is significant, the red tail of the neutral absorption band can overlap considerably with the blue tail of the anionic absorption (as in Ca^2+^-free R-GECO1, see Figure 1D). Therefore, absorption of both the neutral and anionic forms would contribute to the optical density (OD) at a typical excitation wavelength used to excite the anionic form and measure its full fluorescence spectrum for the ϕ. We developed a new method that accounts for this (see Materials and Methods).

Finding the ε for solely the excitable form is also a more involved process due to the presence of two forms of the chromophore. With the alkaline denaturation method, both forms of the chromophore give rise to the denatured form with a known ε. Because of this, the extinction coefficients listed in the original red GECIs papers (11, 14, 15, 19, 21, 22) are effective values and technically equal to a product of ε_e_ and ϱ_e_. We measured the ε for both the neutral and the anionic form of each protein in both Ca^2+^ states (see Materials and Methods).

The first two factors, ϱ_e_ and ϕ_e_, are the same for both one-photon and two-photon excitation. However, the ratios ε_e,sat_ /ε_e,free_ and σ_2,e,sat_ /σ_2,e,free_ might not be equal since ε and σ_2_ can depend differently on the local environment of the chromophore (25–27). Therefore, the overall change in fluorescence could depend on the mode of excitation.

## Materials and Methods

### Protein purification

REX-GECO1 and jRGECO1a were cloned into the pUE backbone via ligation-independent cloning (In-Fusion, Clontech, Mountain View, CA). CAR-GECO1, K-GECO1, R-GECO1, R-GECO1.2, jRCaMP1a, and O-GECO1 were cloned into the pBAD backbone (Thermo Fisher Scientific, Waltham, MA) with restriction cloning. jREX-GECO1 was cloned into the modified pBAD backbone pTorPE with restriction cloning. All plasmids are available in Addgene. They were transformed into DH10B *E. coli* and grown at 30 °C for 2 days in Terrific Broth or Circlegrow (MP Biomedicals, Santa Ana, CA). To induce protein expression in the pBAD plasmids, 0.1% arabinose was added at ~ 5 hours. Bacteria pellets were lysed with BugBuster (EMD Millipore Corp., Burlington, MA) and the lysate was purified with His60 Ni Superflow Resin (Takara Bio, Mountain View, CA). Proteins were buffer-exchanged into pH 7.2 buffers containing 30 mM MOPS, 100 mM KCl, and 10 mM EGTA with or without 10 mM CaCl_2_. Protein characterization was done in these respective buffers, except where noted. Purified proteins were stored short term (≤2 weeks) at 4 °C and long term at −80 °C. The Ca^2+^-free form of K-GECO1 exhibits photoswitching in room light; so the protein sample was stored covered in foil, and all of the photophysical measurements were done with minimal exposure to room light.

### Fluorescence quantum yields

Fluorescence quantum yields (ϕ) were measured with an integrating sphere (Quantaurus-QY; Hamamatsu Photonics, Hamamatsu City, Japan). The solvent-only reference measurement was done with the same cuvette as the sample. Values presented in Table 1 are averages of 3-20 measurements.

With an integrating sphere, it is trivial to measure the ϕ of a sample with only one fluorescent chromophore. However, in these Ca^2+^ sensors, there are two forms of the red chromophore: the neutral, absorbing at ~ 450 nm, and the anionic, absorbing at ~ 570 nm. We measured the ϕ of the excitable form of the chromophore, which could be neutral or anionic. If the relative fraction of the neutral form is significant and the ϕ of the anionic form is needed, then due to spectral overlap both forms could contribute to the optical density (OD) at a typical excitation wavelength used to excite the anionic form and measure its full fluorescence spectrum. We developed the following method to correct for this and applied it to measure the ϕ of the Ca^2+^-free state of jRGECO1a, K-GECO1, jRCaMP1a, CAR-GECO1, O-GECO1, R-GECO1, and R-GECO1.2.

The method requires measurements at two different wavelengths. One is at the typical excitation wavelength, which we call λ_*blue*_, where the neutral form still absorbs, and the other is at the peak of the anionic form absorption, λ_*red*_, where the neutral form absorption becomes insignificant in comparison. With information from these two measurements, and the following equations, we can calculate the ϕ of the anionic form.

In an integrating sphere like the Quantaurus-QY which we used for our measurements, the ϕ is calculated as the integral of the emission spectrum of the sample, *P*, divided by the difference of the integrals of the excitation light intensity transmitted by the reference (solvent only), *R*, and the sample, *S* (Eq. 5). The integral *P* is proportional to the total number of photons emitted by the sample, and *R - S* is proportional to the number of photons absorbed.

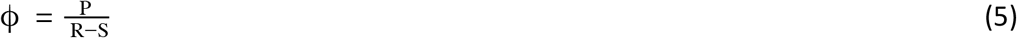

Each integral is written explicitly in Eqs 6, 7, and 8, where λ_ex_ is the excitation wavelength, Δλ is the excitation half-bandwidth (~ 10 nm), *f(*λ) is the fluorescence spectrum, and *R(*λ*) and S(*λ) are the spectral distribution of excitation light intensity reaching the detector.

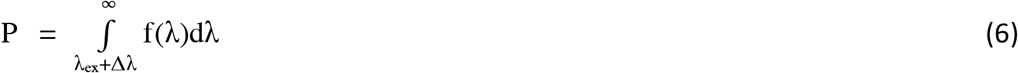

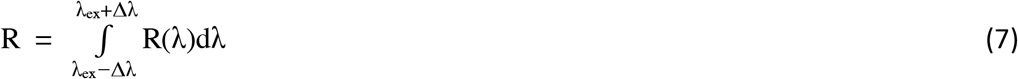

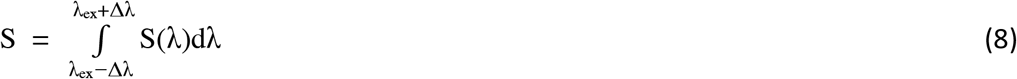

When exciting the sample with λ_*red*_ excitation, the absorbance should belong solely to the anionic form. However, the measured effective ϕ can be either underestimated or overestimated compared to the correct ϕ due to two counteracting factors. First, only part of the emission spectrum can be integrated because λ_*red*_ falls within its blue part. Second, the part of the fluorescence resonant with the excitation light contributes to the integral of the excitation light intensity of the sample *S*, resulting in the underestimation of absorbed photons. This effective, measured, ϕ is defined as *Q* and described by Eq. 9.

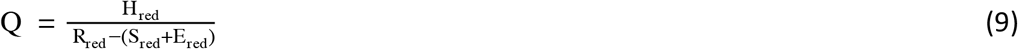

Here, *H_red_* is the sample emission spectrum integrated over the limited wavelength range, which we call the “short” fluorescence integral (Eq. 10). *E_red_* is the sample emission integrated over the bandwidth of excitation, which we call the “very short” integral (Eq. 11). The correct ϕ, *C*, requires the “full” integral, or the integral of the full fluorescence spectrum, *F_red_*, and is described by Eq. 12.

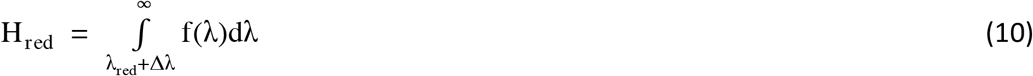

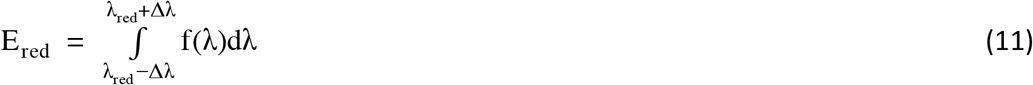

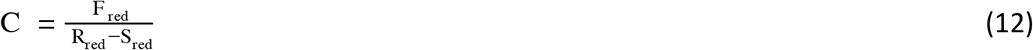

To find the correct ϕ from the effective *Q* value, we measure the fluorescence spectrum with excitation at λ_*blue*_, which is out of the range of emission. With this, we can get the relative values of the full, short, and very short fluorescence integrals. We use the same integration limits as in the λ_*red*_ measurement to get the short integral *H_blue_* and very short integral *E_blue_*. Each set of integration limits are illustrated in Figure S1A in the Supporting Material. Now, we can use the relations *F_red_*/*H_red_* = *F_blue_*/*H_blue_* and *E_red_*/*H_red_* = *E_blue_*/*H_blue_*, and Eq. 9, to find *C* (Eq. 13).

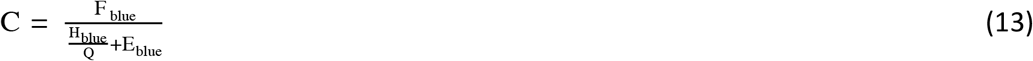

We checked this method with Cresyl Violet in ethanol (fluorescence spectrum ranges from 580-800 nm) with excitation wavelengths λ_*red*_ = 600 nm, 613 nm, and 625 nm and λ_*blue*_ = 565 nm (Figure S1B). Since there is just one chromophore present, the ϕ obtained with λ_*blue*_ and the full fluorescence integral is correct. After applying the method described above, the quantum yields obtained with each λ_*red*_ matched the correct ϕ with errors less than 1%.

### Chromophore pK_a_ values and extinction coefficients

The Ca^2+^-free and Ca^2+^-saturated states of each protein were subjected to stepwise alkaline and acid titrations by gradually adding 2-10 μl of 0.1-1 M NaOH or HCl, respectively, to a sample of 500-1200 μl. Absorption was measured at each step with a LAMBDA 950 Spectrophotometer (PerkinElmer, Waltham, MA), and pH was measured with an Orion™ PeropHecT™ ROSS™ combination pH microelectrode (Thermo Fisher Scientific, Waltham, MA). For the alkaline titrations of the Ca^2+^-saturated state, the protein was first diluted 1:100 into a pH 7.2 buffer with 30 mM MOPS, 100 mM KCl, 50 μM CaCl_2_, and no EGTA, because at a higher pH the *K_d_* of EGTA for Ca^2+^ decreases dramatically (28). To determine the chromophore pK_a_ values, the OD at the peak of the anionic absorbance (even if it shifted throughout the titration) was plotted versus pH in a range where it was not denaturing and the protein was not precipitating (which happened in some instances at the end of the acid titrations). The plot was fit with 1, 2, or 3 pK_a_ values using OriginPro 2017 (OriginLab, Northampton, MA) and Eq. 14 where ODmax_*n*_ is the maximum OD of specie *n* for pK_a,n_ (and was a free parameter).

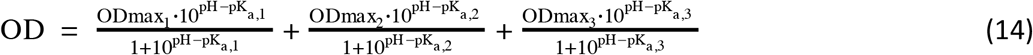

For the titrations of the Ca^2+^-saturated state of jREX-GECO1 and REX-GECO1, the OD at the peak of the anionic form absorbance had to be corrected for absorbance of the neutral form at that position with Eq. 15:

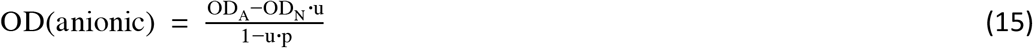

OD_A_ is the OD at the anionic peak position, OD_N_ is the OD at the neutral peak position, *u* is the fraction of neutral absorption at the anionic peak position, and *p* is the fraction of anionic absorption at the neutral peak position. The fraction *u* was determined based on the spectrum at 7.2, and the fraction *p* was based on the spectrum at the pH where the anionic form reached its maximum.

The anionic and neutral forms of the chromophore have distinct extinction coefficients (ε), and we employed the following methods to find each one. To get the εvalues of the anionic form for R-GECO1, R-GECO1.2, O-GECO1, jRGECO1a, REX-GECO1, and jREX-GECO1, we took the OD of the anionic peak at the pH where it reached its maximum during the alkaline titration, and ratioed it with the maximum OD of the alkaline denatured form with a known ε (44,100 M^−1^cm^−1^ (29)). We considered the presence of the immature green chromophore to be insignificant in these proteins. For R-GECO1, R-GECO1.2, O-GECO1, and jRGECO1a, where the anionic peak shifted red at pH ≳ 8.5 in the Ca^2+^-saturated state, we took the maximum OD before the shift. The shift is probably due to a titration of residue(s) near the chromophore and our method provided the ε of the chromophore in the environment corresponding to pH 7.2. To get the ε of the neutral form using the now-known εof the anionic form, we plotted the anionic peak OD versus the neutral peak OD across the titration, and where the dependence was linear, we used the slope to calculate the neutral ε. The OD at the neutral peak was corrected for any contribution of the blue tail of the anionic absorption, using the spectrum at the pH when the anionic form is maximum as a reference. This slope method is further detailed in (24).

For CAR-GECO1, K-GECO1, and jRCaMP1a, the slope method was used to find both the ε of the anionic form and the neutral form because of the significant presence of the immature green chromophore in CAR-GECO1 and the appearance of a 380 nm peak in K-GECO1 and jRCaMP1a during denaturation. To obtain the anionic ε for these proteins, we plotted the OD of the anionic form versus the OD of the ~ 450 nm denatured form and fit the slope at the end of the titration where it was linear and, in the case of K-GECO1 and jRCaMP1a, where the 380 nm peak had already reached its maximum.

### Determination of the relative fraction of the excitable form of the chromophore

The relative fraction of the excitable form of the chromophore in the Ca^2+^-free and Ca^2+^-saturated states for each protein, ϱ_e_, was calculated with Eq. 16 using the absorption spectra collected at pH 7.2.

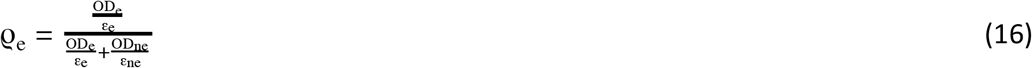

OD is the optical density and ε is the extinction coefficient; the subscripts e and ne stand for excitable and non-excitable form, respectively.

In the case of K-GECO1 and CAR-GECO1, the absorption spectra at pH 7.2 include significant absorption from other, non-red chromophore(s) which overlap with the absorption of the neutral form of the red chromophore. Their absorption spectra were deconvolved using excitation spectra in order to find the OD of the red neutral chromophore before applying the above equation.

For both the Ca^2+^-free and Ca^2+^-saturated spectra of K-GECO1, we fit the red anionic form excitation spectrum (shifted to match the peak position if necessary) to the absorption spectrum and subtracted it, leaving only the absorption of the neutral chromophore and the unknown form (peaking near 400 nm). To this we fit the red neutral form excitation spectrum and took the peak value as the OD of the neutral chromophore. The anionic form excitation spectrum was measured with the Ca^2+^-saturated sample with emission registration at 700 nm. The neutral form excitation spectrum was measured with the Ca^2+^-free sample with emission registration at 530 nm.

For the Ca^2+^-saturated state of CAR-GECO1, we subtracted the excitation spectrum of the immature anionic green chromophore from the absorption spectrum and then fit the red neutral form excitation spectrum under the difference spectrum to find the OD. For the Ca^2+^-free state, we simply fit the red neutral form excitation spectrum to the unadulterated spectrum. The anionic green chromophore excitation spectrum was measured using Ca^2+^-saturated CAR-GECO1 with emission registration at 540 nm. The red neutral form excitation spectrum was measured scanning the Ca^2+^-free sample with emission registration at 540 nm.

### Two-photon characterization

To collect the two-photon absorption spectra, the tunable femtosecond laser InSight DeepSee (Spectra-Physics, Santa Clara, CA) was used to excite the fluorescence of the sample contained in a 3 mm glass cuvette (Starna Cells, Atascadero, CA) within a PC1 Spectrofluorimeter (ISS, Champaign, IL). The laser was automatically stepped to each wavelength over the spectral range with a custom LabVIEW program (National Instruments, Austin, TX), with at least 30 sec at each wavelength to stabilize. We measured two samples per laser scan by using both the sample and reference holders and switching between them with the auto-switching mechanism on the PC1. The laser was focused on the sample through a 45-mm NIR achromatic lens, antireflection coating 750 - 1550 nm, (Edmund Optics, Barrington, NJ). Fluorescence was collected from the first 0.7 mm of the sample at 90° with the standard PC1 collection optics through a combination of either 680/SP, 633/SP, or 694/SP filters together with a 745/SP filter (Semrock, Rochester, NY) to remove all laser scattered light, and a 570/LP filter (Edmund Optics, Barrington, NJ) to remove the residual green fluorescence of the immature chromophore. To correct for wavelength-to-wavelength variations of laser parameters, we used LDS798 (Exciton, Lockbourne, OH) in 1:2 CHCl_3_:CDCl_3_ as a reference standard between 900-1240 nm (30), and Coumarin 540A (Exciton, Lockbourne, OH) in 1:10 DMSO:deuterated DMSO between 700-952 nm (31). Adding the deuterated solvents (Millipore-Sigma, Darmstadt, Germany) was necessary to decrease NIR solvent absorption. The standards were measured at least twice a week in both the sample and the reference holder. All the dyes were magnetically stirred throughout the measurements. Quadratic power dependence of fluorescence in the proteins and standards was checked at several wavelengths across the spectrum.

For the two-photon cross sections, we measured each protein versus Rhodamine 6G in MeOH at 1052 nm (two-photon cross section, σ_2_ = 9.9 GM (32)). Rhodamine 6G was chosen because the value for σ_2_ at this wavelength agrees across multiple literature references ((31) and as discussed in (32)). (Rhodamine B, which was the standard for previous two-photon measurements of some red GECIs, is sensitive to pH with variable σ_2_ values.) Power dependence of fluorescence was recorded with the PC1 monochromator at a wavelength where the OD was ≤ 0.05 and fitted to a parabola with the curvature coefficient proportional to σ_2_. These coefficients were normalized for the differential fluorescence quantum yield at the registration wavelength and the concentrations. The differential quantum yields of the standard and the sample were obtained using the data from the fluorescence quantum yield measurements and selecting an integral range from ~ 5 nm to the left and right of the registration wavelength. Concentrations were determined by Beer’s Law. For Rhodamine 6G in MeOH, ε = 100,500 M^−1^cm^−1^ (32). For most of the proteins (except for the Ca^2+^-saturated state of REX-GECO1 and jREX-GECO1), the σ_2_ at 1052 nm corresponds to the anionic form of the chromophore, and the concentration was determined with the maximum OD of the anionic form and its ε. For the Ca^2+^-saturated state of REX-GECO1 and jREX-GECO1, the OD at the peak of the absorption spectrum and the ε of the neutral chromophore was used to find the concentration.

### Direct measurements of Ca^2+^-saturated/Ca^2+^-free F_1_ and F_2_ ratios

We measured the Ca^2+^-saturated/Ca^2+^-free F_1_ and F_2_ ratios for each protein directly by comparing the Ca^2+^-saturated and Ca^2+^-free states with known concentrations of protein relative to each other. To make the samples, a portion of Ca^2+^-free protein was diluted 10 times to a total volume of 150 μl into a Ca^2+^-saturated buffer (30 mM MOPS, 100 mM KCl, and 10 mM CaCl_2_), and then the pH was adjusted back to 7.2 with 3-5 μl of 0.1 M NaOH. An equivalent volume of 18.2 MΩ water was added to the Ca^2+^-free sample to maintain a known concentration ratio between the two samples.

For the F_2_ ratio spectra, the shape was calculated by dividing the Ca^2+^-saturated two-photon excitation spectrum by the Ca^2+^-free two-photon excitation spectrum. Then, the ratio of the fluorescence signals was measured at three laser wavelengths around the maximum ratio. The setup from the two-photon characterization section was employed, and fluorescence was collected through 745/SP and 770/SP filters (Semrock, Rochester, NY). After applying the known concentration ratio, the three fluorescence signal ratios were normalized to the F_2_ ratio spectrum and the results were averaged.

For the F_1_ ratios, the samples were diluted further into 1 cm glass cuvettes for a total volume of 2 ml. The excitation spectrum for each sample was scanned with the LS 55 Fluorescence Spectrometer (PerkinElmer, Waltham, MA), and the entire fluorescence was collected by setting the emission grating to the 0-th diffraction order. The background fluorescence spectrum of water in a 1 cm glass cuvette was subtracted from each spectrum so that the fluorescence at 650 nm was approximately zero. The ratio between the Ca^2+^-saturated and Ca^2+^-free spectra was calculated and adjusted to account for the concentration difference between the two.

### Emission spectra

The LS 55 Fluorescence Spectrometer (PerkinElmer, Waltham, MA) was used to collect emission spectra. The emission spectra were corrected for spectral sensitivity of the detection system by applying a correction function created with a Spectral Fluorescence Standard Kit (Millipore-Sigma, Darmstadt, Germany).

### Structural analysis

The illustration of the K-GECO1 crystal structure in Figure 1A and structural analyses from the discussion were performed with the UCSF Chimera package (University of California, San Francisco, CA). Chimera is developed by the UCSF Resource for Biocomputing, Visualization, and Informatics (supported by NIGMS P41-GM103311).

### Development and K_d_ determination of jREX-GECO1

To increase the Ca^2+^ affinity of REX-GECO1 (15), the Q306D and M339F mutations were introduced using the QuikChange Mutagenesis Kit (Agilent Technologies, Santa Clara, CA). These two mutations were reported to be responsible for the increased Ca^2+^ affinity of jRGECO1a (19) relative to R-GECO1. The resulting variant was designated as jREX-GECO1. To determine the *K_d_* of jREX-GECO1, Ca^2+^ titrations were performed using EGTA-buffered Ca^2+^ solutions as previously described (11). Briefly, purified proteins were diluted into a series of buffers with free Ca^2+^ concentrations ranging from 0 nM to 39 μM at 25 °C. These buffered Ca^2+^ solutions were prepared by mixing a CaEGTA buffer (30 mM MOPS, 100 mM KCl, 10 mM EGTA, 10 mM CaCl_2_) and an EGTA buffer (30 mM MOPS, 100 mM KCl, 10 mM EGTA) at appropriate ratios. Fluorescence intensities were measured and plotted against Ca^2+^ concentrations and fitted by a sigmoidal binding function to extract the Hill coefficient and *K_d_* for Ca^2+^.

## Results

We studied R-GECO1, R-GECO1.2, CAR-GECO1, O-GECO1, REX-GECO1, jRGECO1a, jRCaMP1a, and K-GECO1. While the chromophores in O-GECO1 and jRCaMP1a have slightly different structures than the red FP chromophore displayed in Figure 1C (Figure S2), we can apply the same analysis because they still undergo the protonation - deprotonation reaction. We also present a new red GECI, jREX-GECO1, which is second generation of REX-GECO1 and incorporates mutations present in jRGECO1a.

The first clue to the mechanism behind the Ca^2+^-dependent change in fluorescence for these red GECIs comes from the one-photon absorption spectra of the Ca^2+^-free and Ca^2+^-saturated states (Figure 2). In all but two of them, the concentration of the anionic form appears to increase from the Ca^2+^-free to the Ca^2+^-saturated state. For jREX-GECO1 (Figure 2C) and REX-GECO1 (Figure 2F), the chromophore distribution appears to shift from mostly anionic in the Ca^2+^-free state to mostly neutral in the Ca^2+^-saturated state. Among those in which the anionic form increases, there is a consistent blue shift of the anionic form absorption peak upon binding to Ca^2+^. The spectral shifts are also apparent in the emission spectra (Figure S3). K-GECO1 (Figure 2B) and jRCaMP1a (Figure 2E) stand out as having both a smaller increase and a shorter blue shift (≤2 nm vs. ≥13 nm) of the anionic form. Based on these differences, these nine red GECIs can be separated into three classes: Class I, which includes jRGECO1a, CAR-GECO1, R-GECO1, R-GECO1.2, and O-GECO1; Class II, which includes K-GECO1 and jRCaMP1a; and Class III, which includes jREX-GECO1 and REX-GECO1. In Classes I and II, the excitable form of the chromophore is anionic regardless of Ca^2+^-binding. In Class III, the excitable form is anionic in the Ca^2+^-free state and neutral in the Ca^2+^-saturated state. Quantitative values of ϱ_e_ are displayed in Table 1.

**Figure 2.**
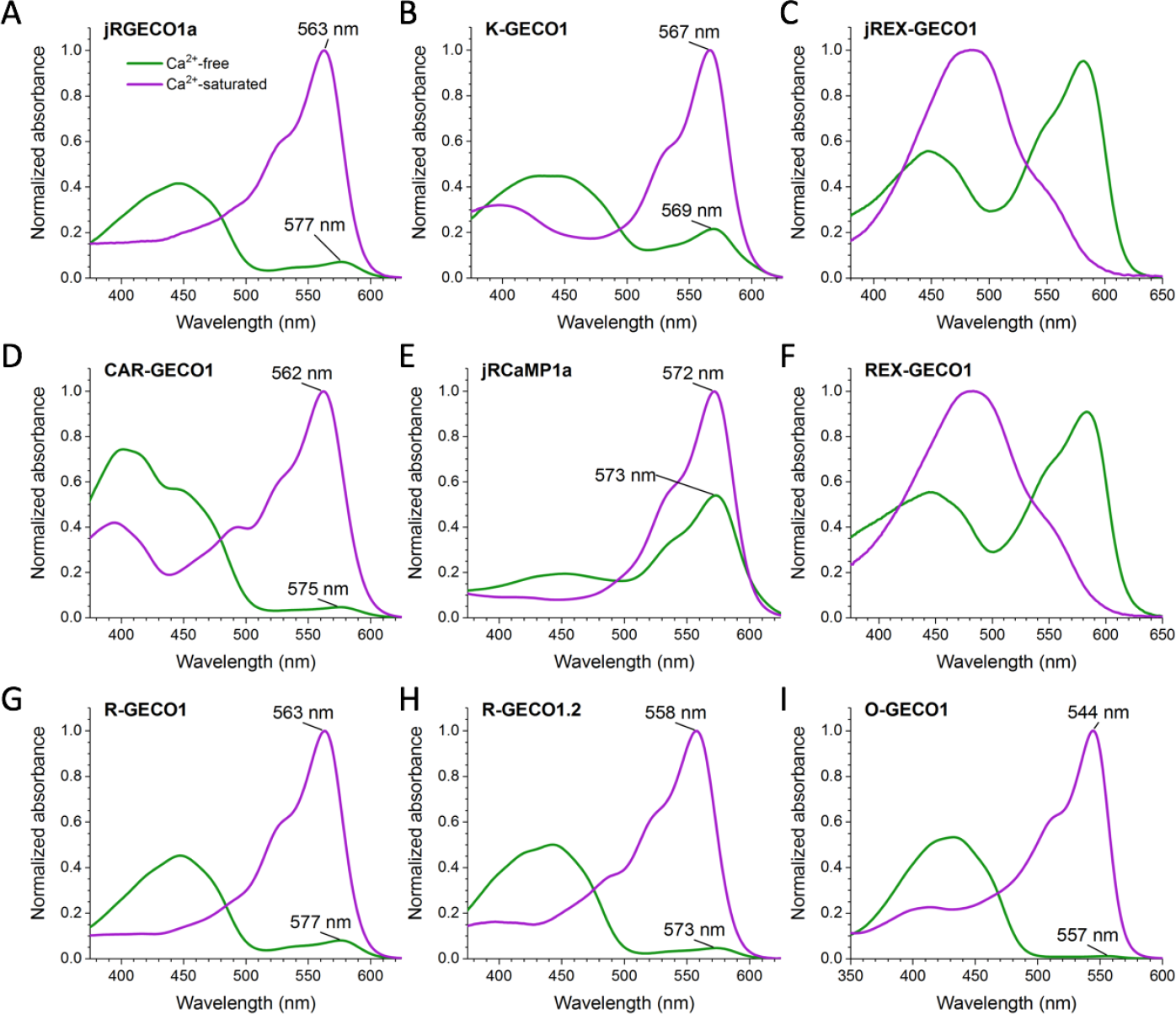
One-photon absorbance spectra of red GECIs in the Ca^2+^-free state (green) and Ca^2+^-saturated state (purple), shown at the same protein concentration in each state and normalized to the peak absorbance in the Ca^2+^-saturated state.

We can gain insight into why ϱ_e_ changes because of Ca^2+^-binding by looking at the absorbance spectra as a function of pH and finding the apparent p*K_a_*s for each Ca^2+^ state (Figure 3, Supplementary Table 1). Starting from Figure 3, we chose to highlight jRGECO1a, K-GECO1, and jREX-GECO1 as examples of their respective classes. Figures for the other six GECIs can be found in the Supplementary Material.

The absorbance pH titrations generally show the neutral form titrating to the anionic form as the pH increases, and they are quite different depending on the Ca^2+^-state. In Class I, the Ca^2+^-saturated states display 2-3 waves of titration, each corresponding to a different p*K_a_* value. This is likely due to titratable residues near the chromophore that affect its p*K_a_* as well as the absorption of the anionic form (16, 33, 34). For Ca^2+^-saturated jRGECO1a, there are three apparent p*K_a_* values (Figure 3A, inset), and the absorption peak of the anionic form shifts to three distinct wavelengths throughout the titration (Figure 3A): 577 nm at ~ pH ≤5.5, 563 nm from ~ pH 6.5-9, and 575 nm at ~ pH ≥9. In the Ca^2+^-free pH titration of jRGECO1a (Figure 3B), the peak position of the anionic form stays at 577 nm, and there is just one p*K_a_* (Figure 3B, inset). The titrations in Class II were best fit with a single p*K_a_*, except for Ca^2+^-free K-GECO1, which displayed a broad titration curve which could only be satisfactorily fit with two p*K_a_* values (Figure 3D, inset). There are subtle spectral shifts of the anionic peak in both Ca^2+^ states of Class II. For Ca^2+^-free K-GECO1, it shifts blue as it increases and red as it denatures. The pH titrations in Class II are also characterized by a peak at about 380 nm that irreversibly appears at alkaline pH, which may be due to the denatured form of the cleaved protein chromophore as shown in Figure S2 (21, 23, 35–37).

Interestingly, in Class III, when titrating the Ca^2+^-saturated state from neutral to acidic pH, the spectrum unexpectedly morphs from the broad ~ 490 nm peak into two peaks that presumably belong to the neutral and anionic forms of the chromophore (Figure 3E). However, there is a clean isosbestic point between the 490 nm form to the 570 nm form when titrating from neutral to alkaline pH (Figure 3F), suggesting that the 490 nm form is singularly neutral. The alkaline titration can be fit with one very high p*K_a_* (11.63 for jREX-GECO1). The Ca^2+^-free titrations for Class III are best fit with three p*K_a_* values (Fig 3G), and the spectral shifts of the anionic form reflect that.

**Figure 3.**
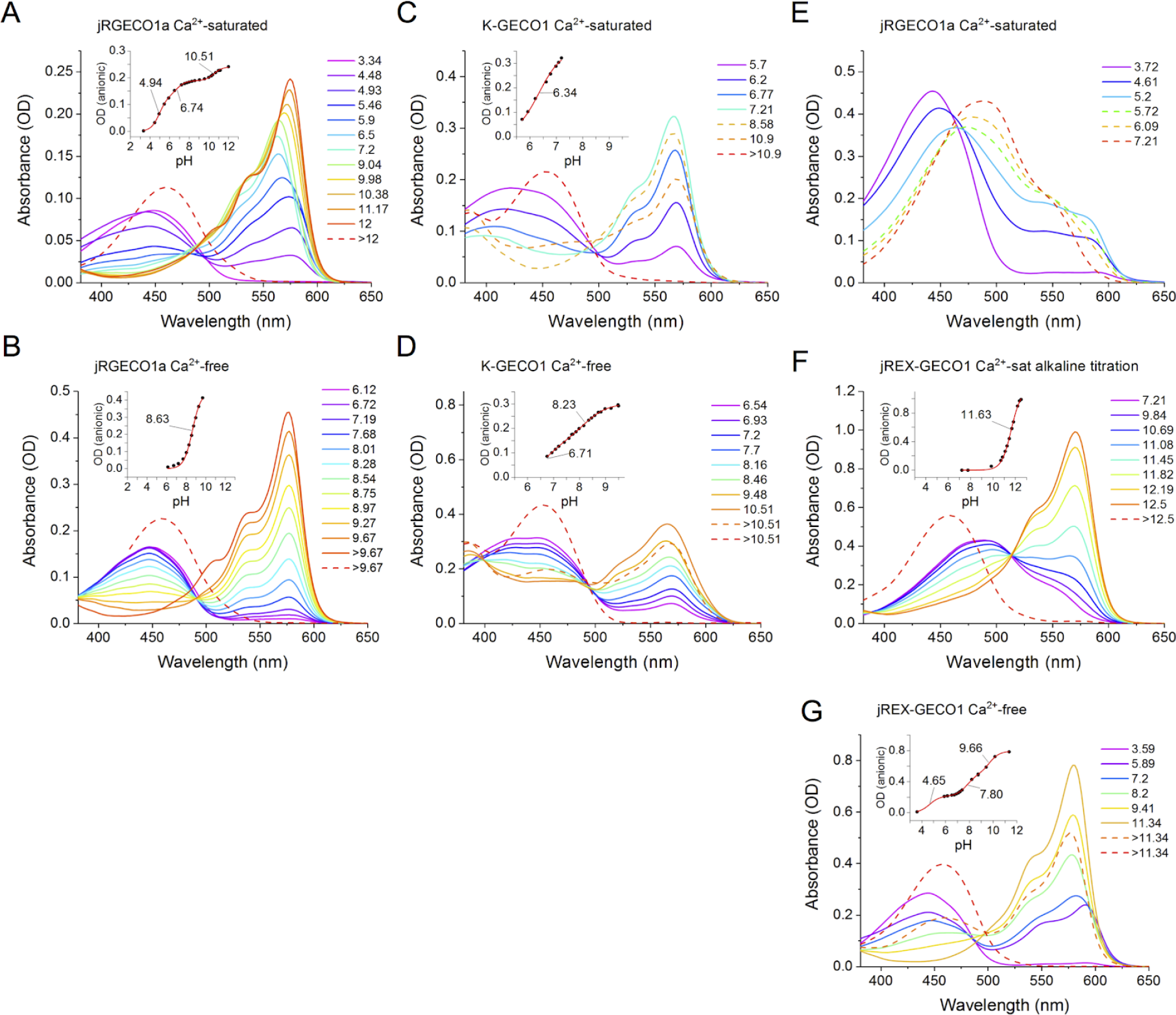
Absorbance pH titrations and p*K_a_* curves (*insets*) for jRGECO1a, K-GECO1, and jREX-GECO1, measured in either Ca^2+^-free or Ca^2+^-saturated buffer. Each spectrum was measured at the pH value designated in the legends. The solid spectra follow the increase of the anionic form peak, and the dashed spectra follow the decrease. Except for (*E*), the final spectrum (red dashed line) belongs to the denatured chromophore. (*F, G*) For clarity, Ca^2+^-saturated jREX-GECO1 pH titration are split into the acid and alkaline titrations, respectively. (*Insets*) Fitted p*K_a_* values are indicated on the curve. Note that for each protein, the Ca^2+^-saturated and Ca^2+^-free plots have an equal pH scale.

For Classes I and II where the excitable form is always anionic, the Ca^2+^-dependent changes in ϱ_A_ qualitatively follow from the shifts in apparent p*K_a_* (Supplementary Table 1). In Class I, if we consider the Ca^2+^-saturated p*K_a_* with the largest amplitude change of the curve, which is always the first (most acidic) one, it is usually about 3.5 pH units smaller than the Ca^2+^-free p*K_a_*. In Class II, for K-GECO1, the p*K_a_* shift is about 1.2 pH units if we take the average of the two apparent p*K_a_*s in the Ca^2+^-free state, and jRCaMP1a has a shift of only 0.3 pH units. Class II exhibits smaller p*K_a_* shifts than Class I, and the Ca^2+^-saturated/Ca^2+^-free ϱ_A_ ratio reflects this: in Class I, the ratio ranges between 14 and 121, while in Class II, the ϱ_A_ ratio is 3 for K-GECO1 and 1.5 for jRCaMP1a.

**Table 1.**
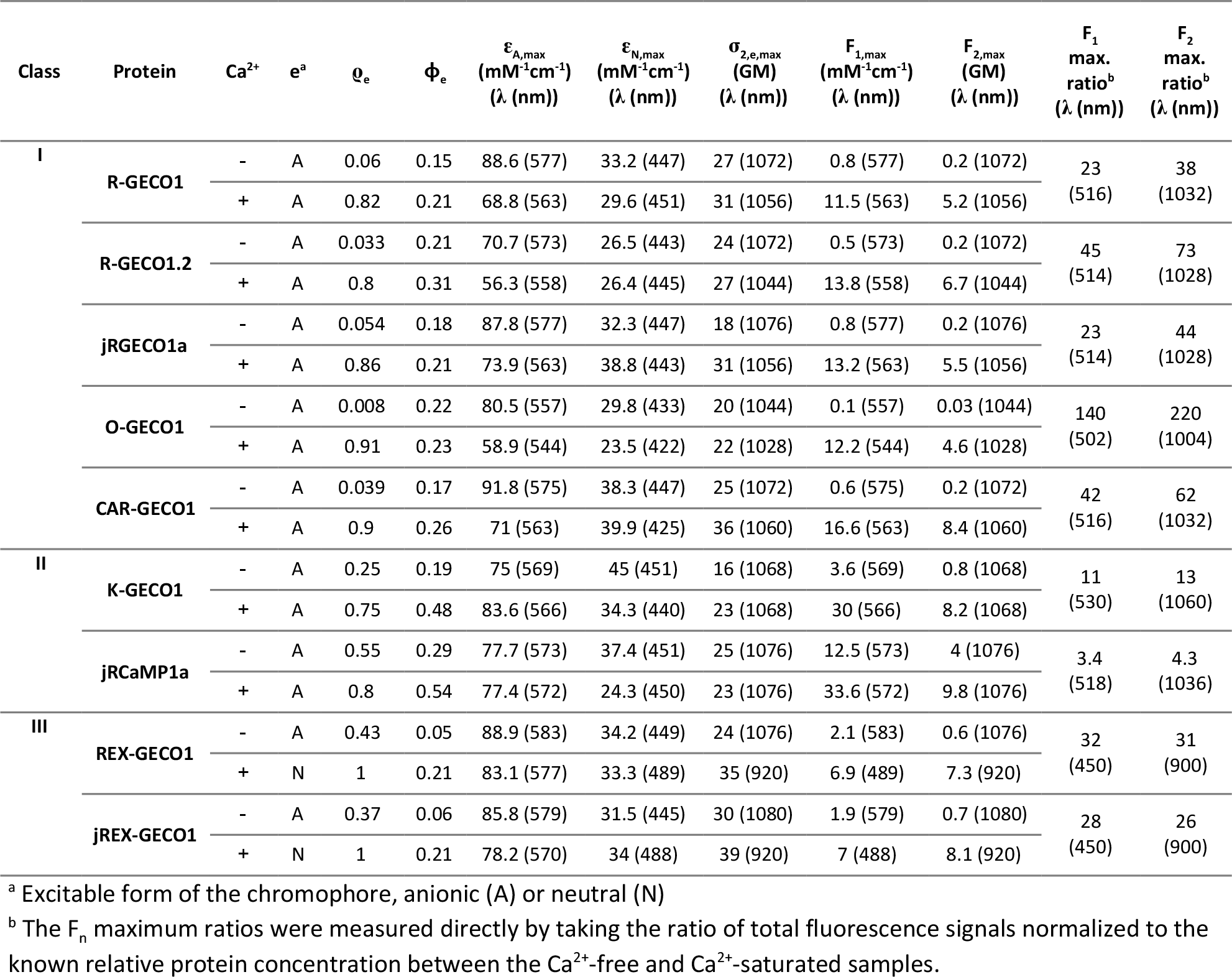
Photophysical properties of the nine red GECIs.

The next clue to the mechanism of the Ca^2+^-dependent change in fluorescence is ϕ_e_ (Table 1). In Class I, ϕ_e_ ranges from 0.15-0.31, and ϕ_e_ increases in the Ca^2+^-saturated state, from an insignificant change in O-GECO1 to 1.5 times in CAR-GECO1. In Class II, the ϕ_e_ values of the Ca^2+^-saturated state are higher than Class I (up to 0.55), and the Ca^2+^-dependent increase of ϕ_e_is more prominent: 1.85 times for jRCaMP1a and 2.5 times for K-GECO1. Class III has the largest increases of ϕ_e_ (2.5 - 3.5 times) because while the Ca^2+^-saturated φs of the neutral form are just 0.21, the φs of the Ca^2+^-free anionic form are much lower (0.05 - 0.06). In every class the ϕ_e_increases upon binding Ca^2+^, which helps to maximize the F_n_ ratios, although the degree to which it does depends on the class.

The final factor contributing to the Ca^2+^-saturated/Ca^2+^-free F_n_ ratios is the one that separates one-photon excitation (1PE) from two-photon excitation (2PE): the molecular absorption coefficient at the excitation wavelength, ε_e_ or σ_2,e_. We directly measured both the F_1_ and F_2_ ratios as a function of wavelength for each protein by taking the ratio of excitation spectra for Ca^2+^-saturated and Ca^2+^-free samples normalized to known relative concentrations of protein. When plotted together versus transition wavelength (corresponding to one-half the laser wavelength in the two-photon case), the shapes of the F_n_ ratios look similar for both modes of excitation (Figure 4A-C). However, the absolute values are not always the same, especially at the excitation wavelengths where the ratios are maximum, which is marked with a vertical dashed line. We call this particular transition wavelength λ_Rmax_. In Classes I and II, the ratio spectra have a double-humped nature because of the spectral shift of the anionic form between Ca^2+^ states, and we selected the shorter wavelength hump as λ_Rmax_ for consistency. For jRGECO1a (Figure 4A), the F_2_ ratio at λ_Rmax_ is about twice as large as the F_1_ ratio. For K-GECO1 (Figure 4B), the F_2_ ratio is still slightly larger, while for jREX-GECO1 (Figure 4C) there is no appreciable difference.

To explain the differences between the F_1_ and F_2_ ratios, Figures 4D-F illustrate the one-and two-photon absorption of the excitable form normalized to ε_e_ and σ_2,e_, respectively. At λ_Rmax_ (vertical dashed line), the σ_2,e_ of jRGECO1a is doubled in the Ca^2+^-saturated state, and because of the spectral shift between the Ca^2+^-states, the ε_e_ is essentially equal for both (Figure 4D). Curiously, the maximum ε_e_ for jRGECO1a actually decreases. K-GECO1 (Figure 4E) displays a smaller increase in the Ca^2+^-saturated σ_2,e_ at λ_Rmax_ (jRCaMP1a has no increase, see Figure S4E) and a very small increase in ε_e_. Finally, jREX-GECO1 (Figure 4F) has relatively equivalent increases of ε_e_ and σ_2,e_ from the Ca^2+^-free to the Ca^2+^-saturated state at λ_Rmax_. Figures 4G-I show the separate F_1_ (dashed lines) and F_2_ (solid lines) brightness spectra for the Ca^2+^-free and Ca^2+^-saturated states. These were determined with independent measurements of ϱ_e_, ϕ_e_, σ_2,e_, and ε_e_ to calculate F_1_ and F_2_ at a wavelength where only the excitable form contributes to the fluorescence. Since jREX-GECO1 is best excited at about 900 nm under 2PE, it is an excellent candidate for two-color two-photon imaging together with a green FP-based probe. This has been demonstrated with its predecessor REX-GECO1 (15).

The final question remains: of the three different factors, which one matters the most? We have found that the answer to this question depends on the indicator class. The Ca^2+^-saturated/Ca^2+^-free ratios of each parameter, ϱ_e_, ϕ_e_, and the molecular absorption coefficients σ_2,e_ and ε_e_ at λ_Rmax_ are visualized in Figure 5. The insets show the total F_n_ ratios, both calculated from the independent measurements of each parameter (tan bars) as well as directly measured (green bars). Representing Class I, jRGECO1a (Figure 5A) has a ϱ_e,sat_/ϱ_e,free_ ratio that far exceeds the other ratios. Again, the change in σ_2,e_ compared to ε_e_ causes the F_2_ ratio to be larger than the F_1_ ratio. From Class II, K-GECO1 (Figure 5B) has comparable contributions to the overall F_n_ ratios from changes in both ϱ_e_ and ϕ_e_. Lastly, in Class III, all three factors play a significant role, as illustrated by the example of jREX-GECO1 (Figure 5C).

**Figure 4.**
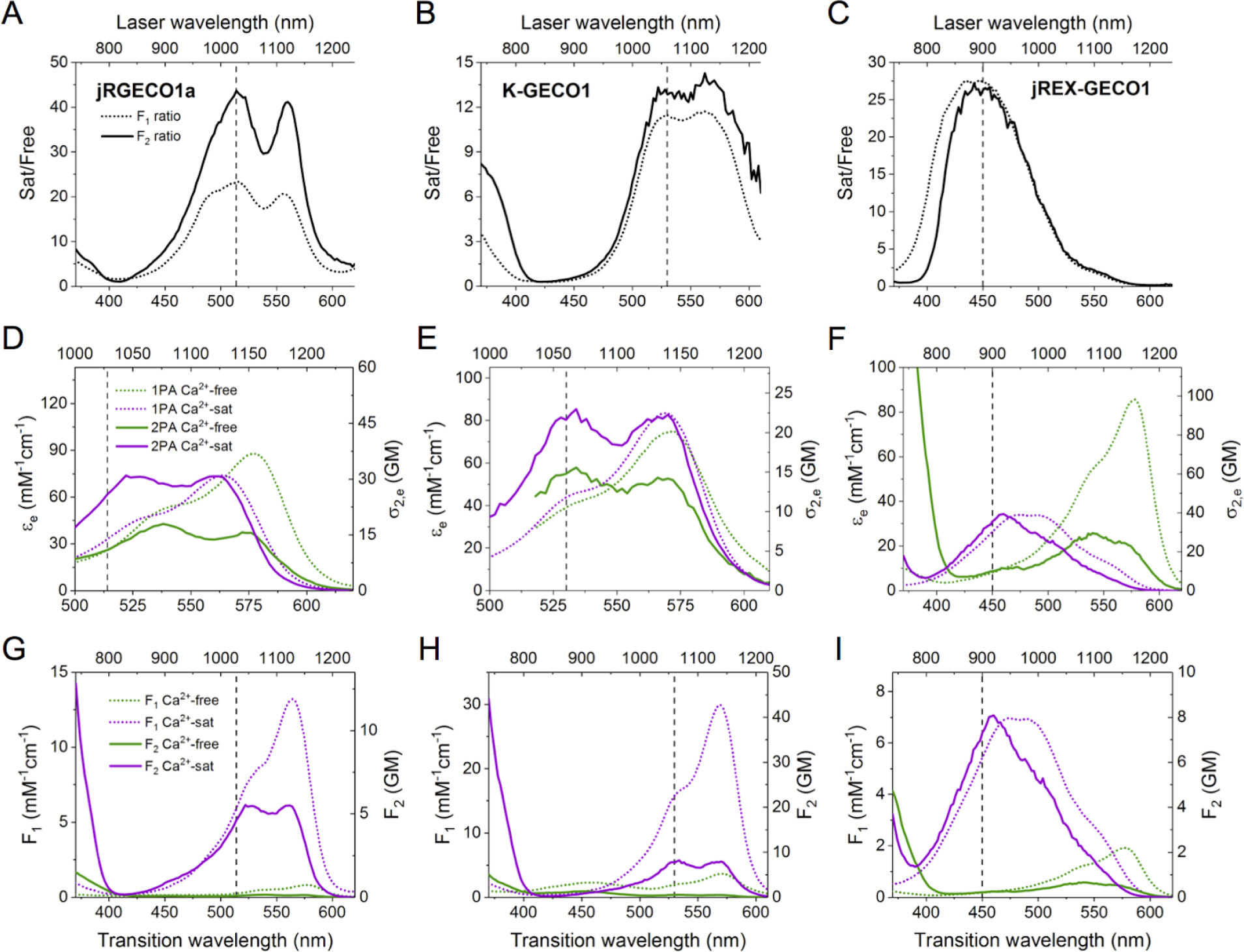
Spectral analysis of jRGECO1a, K-GECO1, and jREX-GECO1, representing Classes I, II, and III of red GECIs, respectively. The vertical dashed line on each plot indicates the transition wavelength of excitation where the Ca^2+^-saturated/Ca^2+^-free F_1_ and F_2_ ratios are approximately maximum (λ_Rmax_). (*A-C*) Ca^2+^-saturated/Ca^2+^-free (Sat/Free) F_1_ and F_2_ ratios as a function of excitation wavelength, measured directly by taking the ratio of the integrated fluorescence signal normalized to the known relative protein concentration between the Ca^2+^-free and Ca^2+^-saturated samples. (*D-F*) One-photon absorption (1PA) and two-photon absorption (2PA) of the excitable form of the chromophore plotted according to ε(left axis) and σ_2_ (right axis), respectively, for the Ca^2+^-free and Ca^2+^-saturated states. (*G-I*) Spectra of the one-photon brightness (F_1_, dotted lines, left and bottom axes) and two-photon brightness (F_2_, solid lines, right and top axes) of Ca^2+^-free and Ca^2+^-saturated states.

**Figure 5.**
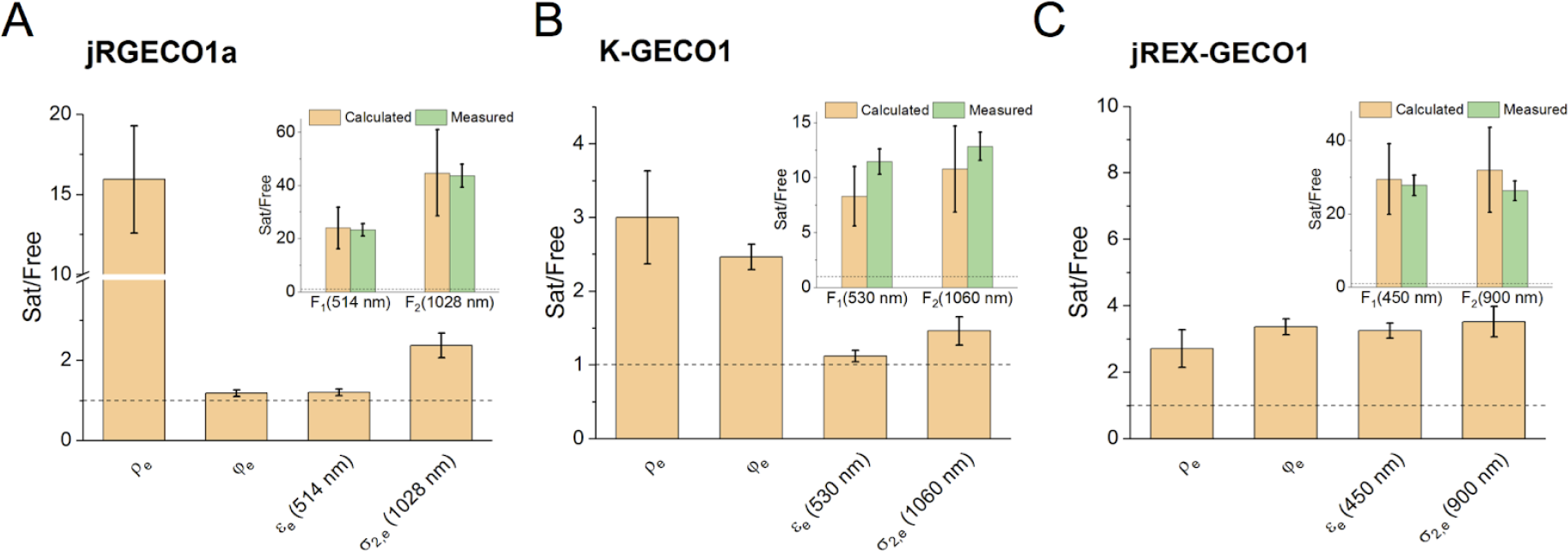
jRGECO1a (*A*), K-GECO1 (*B*), and jREX-GECO1 (*C*) Ca^2+^-saturated/Ca^2+^-free (Sat/Free) ratios for ϱ_e_, ϕ_e_, ε_e_ and σ_2,e_ at λ_Rmax_. Insets: Overall Ca^2+^-saturated/free F1 and F2 ratios, calculated from the independent measurements of ϱ_e_, ϕ_e_, ε_e_, and σ_2,e_ (tan) and directly measured (green). The horizontal dashed line marks a Sat/Free ratio of 1 (no change) for each plot.

## Discussion

We set out to determine the factors that contribute to the Ca^2+^-dependent fluorescence change in nine red GECIs under both one-photon and two-photon excitation. The key parameters that we considered were: 1) the fraction of the protein in the excitable protonation state (ϱ_e_); 2) the quantum yield for the excitable state (ϕ_e_); and 3) the extinction coefficient (ε_e_, for 1PE) or two-photon cross section (σ_2,e_, for 2PE) at the excitation wavelength. We found that the nine proteins could be separated into three different classes, each of which relied on different factors. The fluorescence change in Class I (jRGECO1a, R-GECO1, R-GECO1.2, CAR-GECO1, and O-GECO1) is mainly due to a dramatic increase in the concentration of the anionic form when bound to Ca^2+^, much like GCaMP6m (24). In Class II (K-GECO1 and jRCaMP1a), an increase in ϕ_e_ rivals the increase in ϱ_e_. In Class III (jREX-GECO1 and REX-GECO1), which is a unique class because the excitable form of the chromophore switches from anionic in the Ca^2+^-free state to neutral in the Ca^2+^-saturated state, all three factors contribute nearly evenly to the change in fluorescence. Classes I and II were previously alluded to in (21), where the importance of the change in the chromophore p*K_a_* between Ca^2+^-states was highlighted for R-GECO1, and the change in ϕ was seen to be more important for the RCaMP series (which jRCaMP1a was based on).

The spectral shapes of the Ca^2+^-saturated/Ca^2+^-free fluorescence excitation ratios are very similar under 1PE and 2PE. Interestingly, for Class I, the value of the fluorescence ratio is greater under 2PE compared to 1PE at λ_Rmax_. In Class II, this is also the case for K-GECO1 but not significant for jRCaMP1a. In these two classes the excitable protonation state of the chromophore is always deprotonated/anionic. When the F_2_ ratio is greater than the F_1_ ratio, it is because the σ_2_ of the anionic form increases upon Ca^2+^-binding while the εchanges little or not at all. The increase in σ_2_ can be explained by its higher sensitivity to the local electric field compared to ε (25–27). This sensitivity is due to a factor describing the change of the permanent dipole moment upon excitation that enters the expression for σ_2_, and the electric field affects this factor through polarizability.

Notably, the increase in σ_2_ upon binding Ca^2+^ is always paralleled by a shift of the one-photon absorption peak to a shorter wavelength. We previously developed a physical model ((25–27)) that predicts that both of these changes will happen in the anionic chromophore if the electric field directed from the center of the phenolate ring to the center of the imidazole ring increases. In red GECIs, the chromophore is oriented such that its phenolate oxygen points toward the opening of the barrel at the site of circular permutation where the Ca^2+^-sensing domain is fused to the FP (as observed for RCaMP (PDB ID: 3U0K (21)), R-GECO1 (PDB ID: 4I2Y (21)), and K-GECO1 (PDB ID: 5UKG (22)). Due to this geometry, the conformational change of the protein when it binds Ca^2+^ would likely modulate the electric field along the phenolate-imidazole axis. We propose that for the indicators where σ_2_ of the anionic form increases, the field in this direction increases and positive charges move closer to the phenolate oxygen and/or negative charges move further away.

Upon binding Ca^2+^, the indicators in Class I show the greatest increase in σ_2_ and largest blue shifts (≥13 nm), K-GECO1 shows an intermediate increase in σ_2_ and a 2 nm shift, and jRCaMP1a virtually does not change σ_2_ and has a 1 nm shift. These differences might be due to the fact that the indicators in Class I have a positive Lys78 residue hydrogen-bonded to the phenolate oxygen of the chromophore in the Ca^2+^-saturated state, but in K-GECO1a and jRCaMP1a only neutral residues (Asn32 and Thr294, respectively) and water molecules (W636 and W467, respectively) are bound to that oxygen (21, 22). If those hydrogen bonds are disrupted in the Ca^2+^-free state of Class I indicators and K-GECO1, then this could explain the largest changes in σ_2_ for Class I and smaller changes for K-GECO1. The negligible change in σ_2_ for jRCaMP1a may be because there is no significant redistribution of charges at the opening near the chromophore.

The changes in ϱ_e_ for Class I and Class II qualitatively follow the same trend as the changes in σ_2_, which can be explained on similar electrostatics grounds. The redistribution of charges around the chromophore with the positive charge coming closer to the phenolic oxygen results in the shift of the acid-base equilibrium of the chromophore towards its deprotonated, anionic form. Conversely, the sensors in Class III were initially developed by mutating the positive Lys78 in R-GECO1 to the negative Glu, stabilizing the neutral form in the Ca^2+^-saturated state. In this case an observed moderate increase in σ_2_ at 900 nm of the excitable form (see Table 1) is due to the transition of the chromophore from the anionic to the neutral form.

We also observe that ϕ of the anionic form increased in the Ca^2+^-saturated state in all cases except O-GECO1 (where it did not change) and Class III proteins (where no anionic form is present in the Ca^2+^-saturated state). This effect can also be explained by the same changes in electric field upon binding Ca^2+^ as described above. Quantum chemistry calculations predict that the nonradiative decay of the excited state of the red FP chromophore in vacuum proceeds through the substantial twisting around the phenolate exocyclic C-C bond which is coupled with the electron density transfer from the phenolate to the imidazolinone and acilimine groups (38). If the field increases in the direction from phenolate to imidazolinone (i.e. in the direction of electron transfer), the activation energy for the electron transfer will increase and the nonradiative decay rate will be slowed down. That will result in the increase of the fluorescence quantum yield. The independence of quantum yield on conformational changes in GCaMP6m (24) and in O-GECO1 (Table 1), is probably due to a lack of an acylimine group in their chromophore structure. In fact, it was shown (38) that the electronegativity of acylimine makes the described charge transfer channel of nonradiative relaxation highly probable specifically in red FP chromophore. A detailed physical model and quantitative description of the quantum yield dependence on electric field in red GECIs will be described in our forthcoming publication. It is interesting to note that all three factors, ϱ_e_, ϕ_e_, and σ_2,e_ synergistically increase in almost all the red GECIs due to a favorable geometric arrangement of the chromophore with respect to the fusion site and to the local electrostatic/hydrogen bonding interactions.

In summary, red GECIs can be distinguished into three classes, and the factors contributing to the Ca^2+^-dependent change in fluorescence vary based on the class. In some cases, the change under one-photon excitation was different than the change under two-photon excitation. We suggest that these differences should be carefully considered when further evolving these red GECIs.

## Author contributions

R.S.M. and M.D. designed and conducted research. J.W. and Y.Q. developed jREX-GECO1. Y.S. provided plasmids. R.S.M. analyzed data and wrote the manuscript. R.E.C., T.E.H., and M.D. supervised research. R.S.M., Y.Q., Y.S., R.E.C., T.E.H., and M.D. revised the manuscript.

## Acknowledgments

We thank A. Drobizhev for the custom LabVIEW program, P. Augustin for her technical help, and M. Thomas and E. Johnson for their feedback on the manuscript. This work was supported by grants from NIH (U01 NS094246 and U01 NS090565), NSERC (RGPIN 288338-2010), CIHR (FS 154310), and Brain Canada. R.S. Molina is supported by the NIH NINDS Fellowship under Award Number F31NS108593. She is particularly indebted to NINDS program staff E. Talley and S. Korn for their encouragement and support.

## SupplementaryInformation

**Figure S1.**
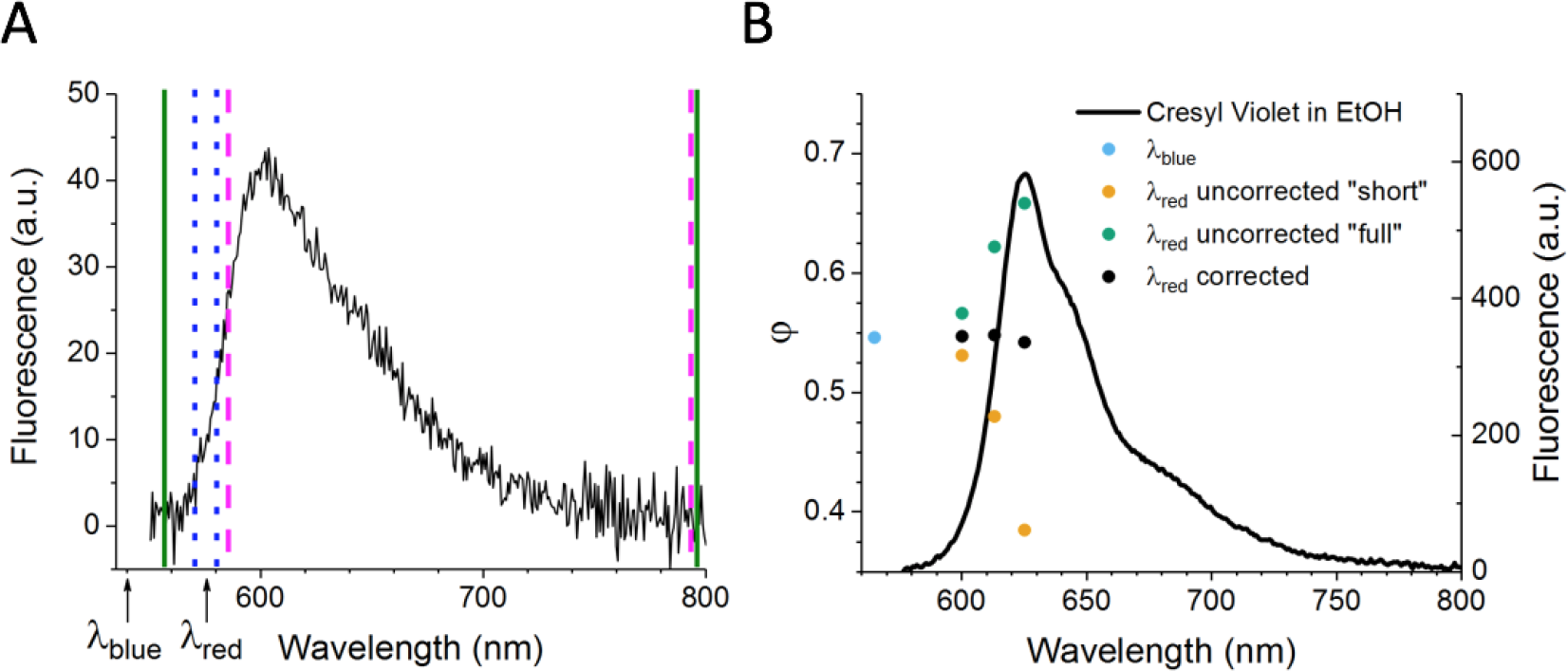
Analytical method to find correct fluorescence quantum yield (ϕ). (*A*) Example fluorescence spectrum (shown is Ca^2+^-free R-GECO1), excited with λ_*blue*_ = 540 nm. Green solid lines indicate the “full” fluorescence integral limits, blue dotted lines the “very short” limits, and magenta dashed lines the “short” limits. λ_*blue*_ and λ_*red*_ are indicated with arrows. (*B*) Quantum yields measured by exciting Cresyl Violet in EtOH at wavelengths resonant with the emission spectrum can be corrected with the method explained in the text. The line corresponds to the emission spectrum of Cresyl Violet in EtOH (right axis). The points represent particular ϕ values (left axis). Blue: the Φ measured at λ_*blue*_, which is the correct φ; Orange: the effective “short” ϕ *Q* at each λ_*red*_; Green: the “full” ϕ values before correcting for the part of the emission *E_red_* included with the sample excitation light intensity *S*; Black: the corrected ϕ *C* for each λ_*red*_.

**Figure S2.**
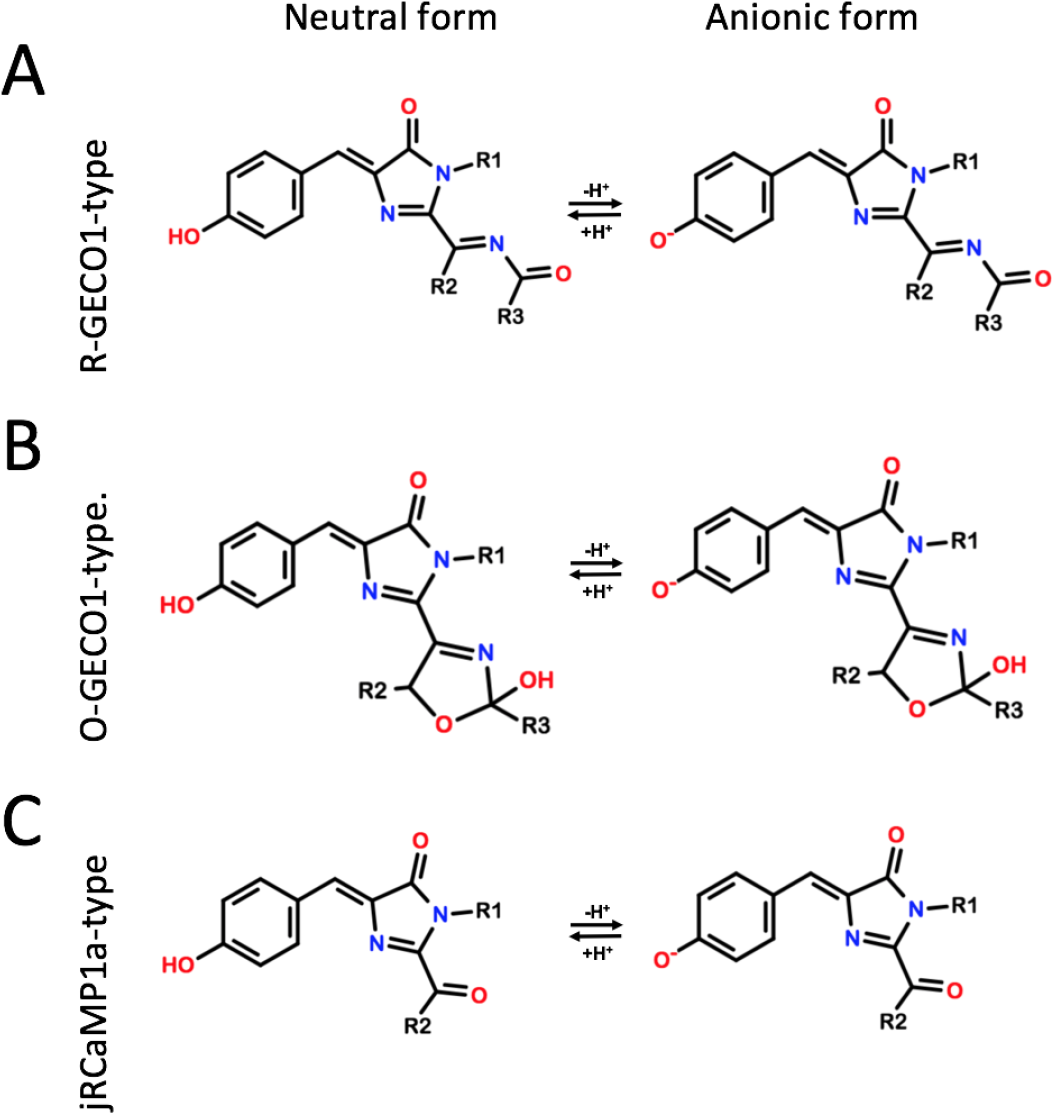
Illustration of the neutral and anionic forms of the chromophores in the red GECIs under study. (*A*) R-GECO1-type chromophore, same as Figure 1C in main text. (*B*) O-GECO1-type chromophore (1). (*C*) jRCaMP1a-type chromophore (2).

**Supplementary Table 1.**
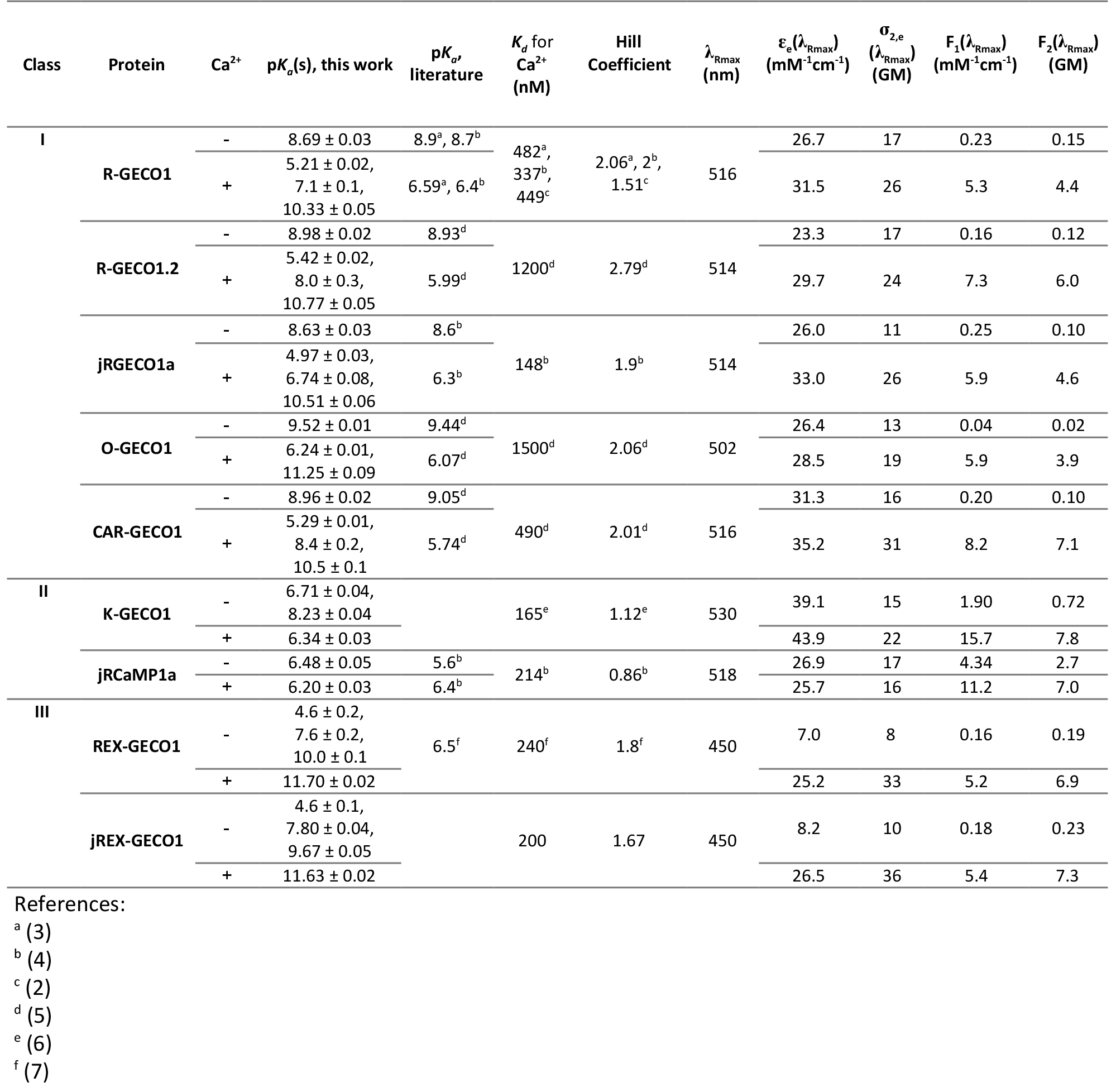
Additional biophysical properties of the red GECIs.

**Figure S3.**
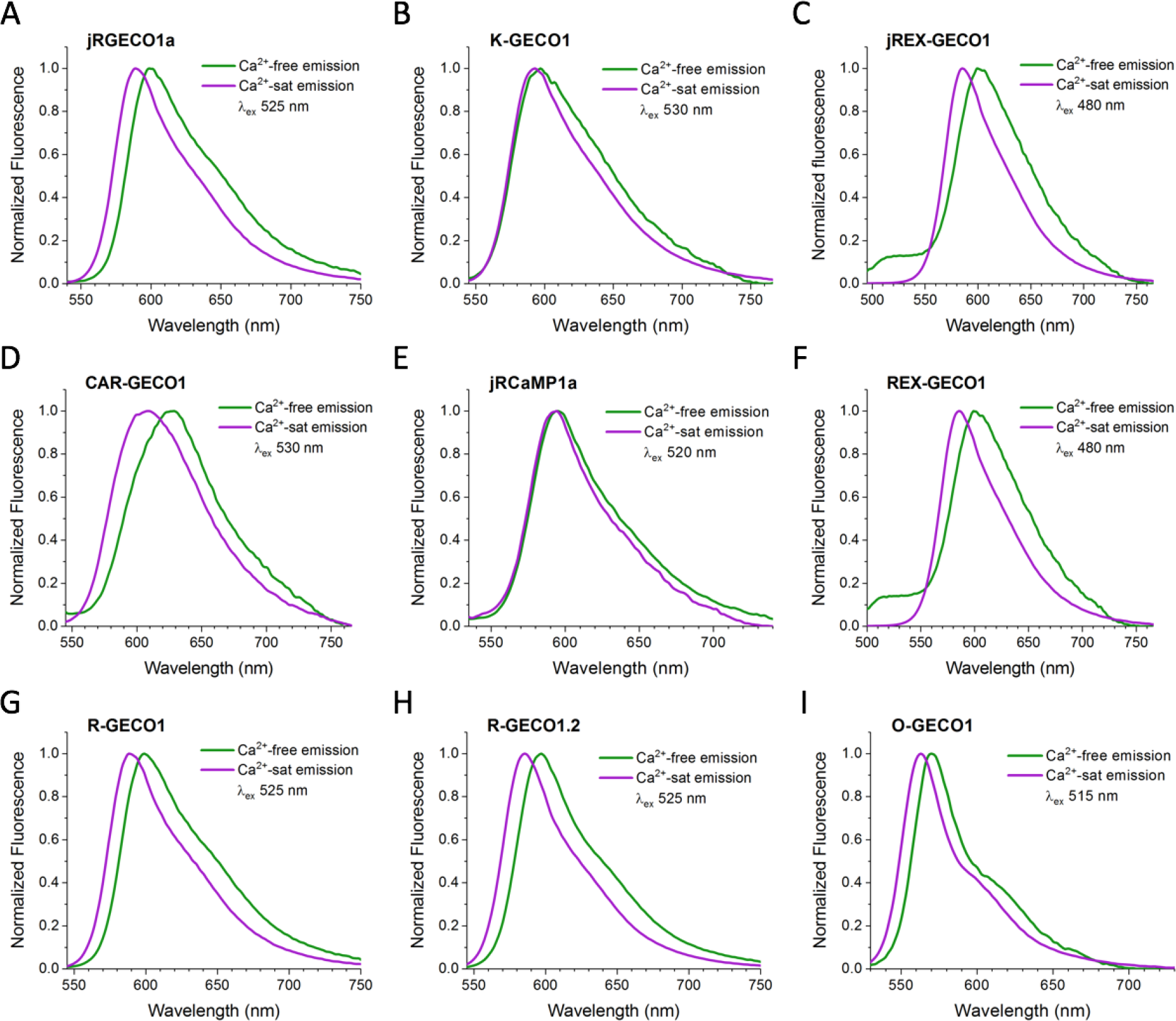
Emission spectra of the Ca^2+^-free and Ca^2+^-saturated states. All are normalized to 1. The excitation wavelength of light used to excite both states of each protein is indicated on each plot.

**Figure S4.**
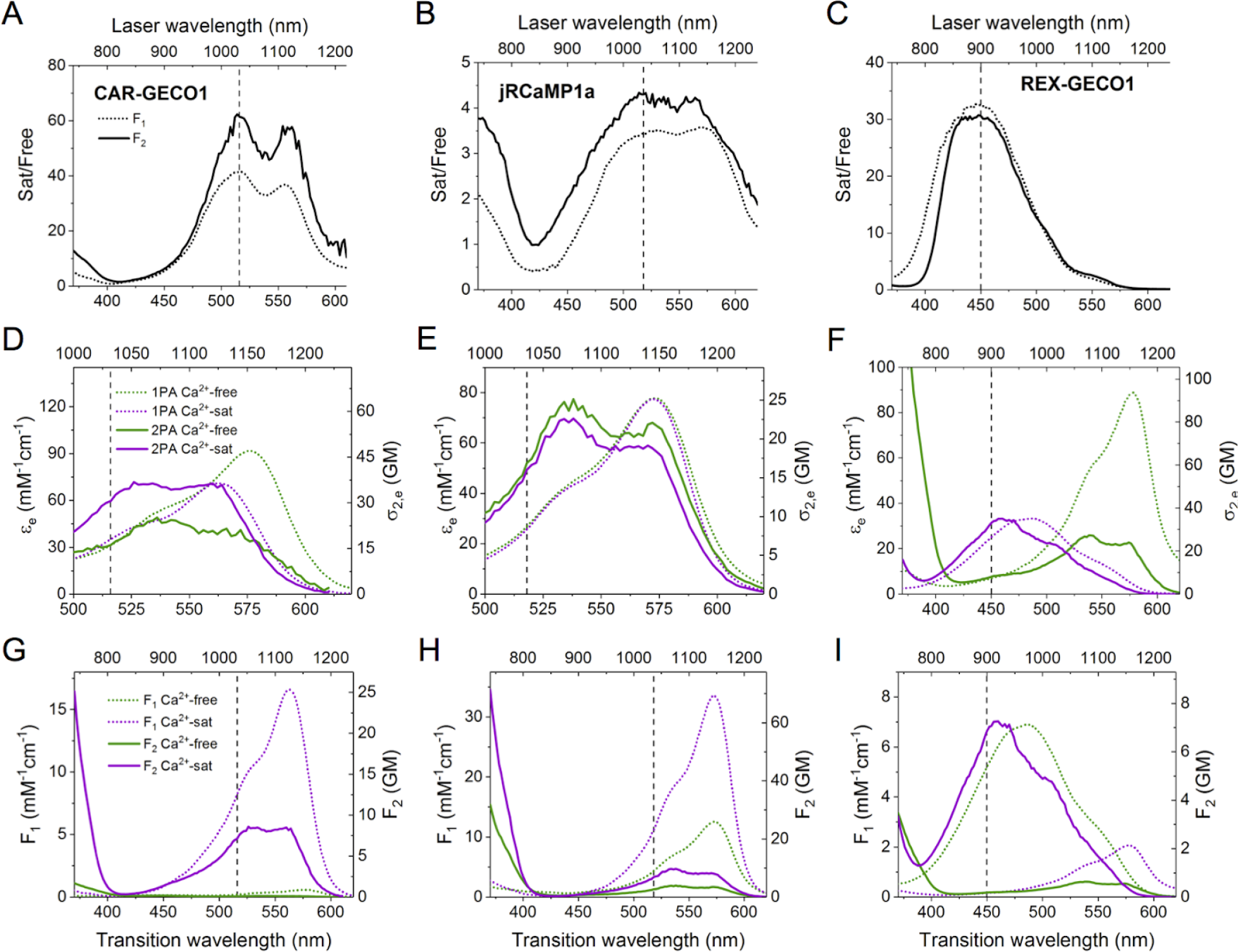
Spectral analysis of CAR-GECO1, jRCaMP1a, and REX-GECO1. Plots are set up as in Figure 4 of main text. The vertical dashed line on each plot indicates the transition wavelength of excitation where the Ca^2+^-saturated/Ca^2+^-free F_1_ and F_2_ ratios are approximately maximum (λ_Rmax_). (*A-C*) Ca^2+^-saturated/Ca^2+^-free (Sat/Free) F_1_ and F_2_ ratios as a function of excitation wavelength, measured directly by taking the ratio of the integrated fluorescence signal normalized to the known relative protein concentration between the Ca^2+^-free and Ca^2+^-saturated samples. (*D-F*) One-photon absorption (1PA) and two-photon absorption (2PA) of the excitable form of the chromophore plotted according to ε(left axis) and σ_2_ (right axis), respectively, for the Ca^2+^-free and Ca^2+^-saturated states. (*G-I*) Spectra of the one-photon brightness (F_1_, dotted lines, left and bottom axes) and two-photon brightness (F_2_, solid lines, right and top axes) of Ca^2+^-free and Ca^2+^-saturated states.

**Figure S5.**
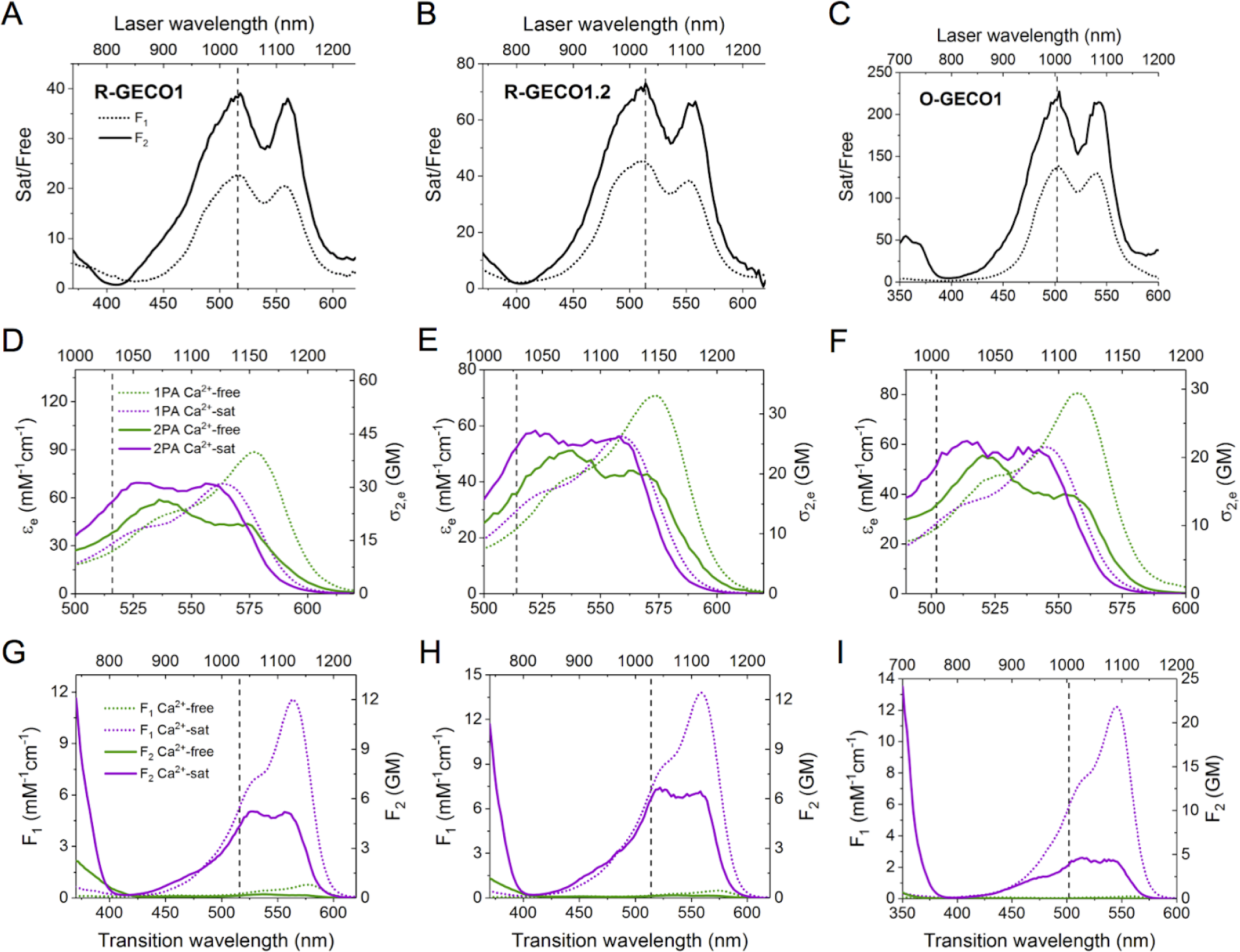
Spectral analysis of R-GECO1, R-GECO1.2, and O-GECO1. Plots are set up as in Figure 4 of main text and Figure S4.

**Figure S6.**
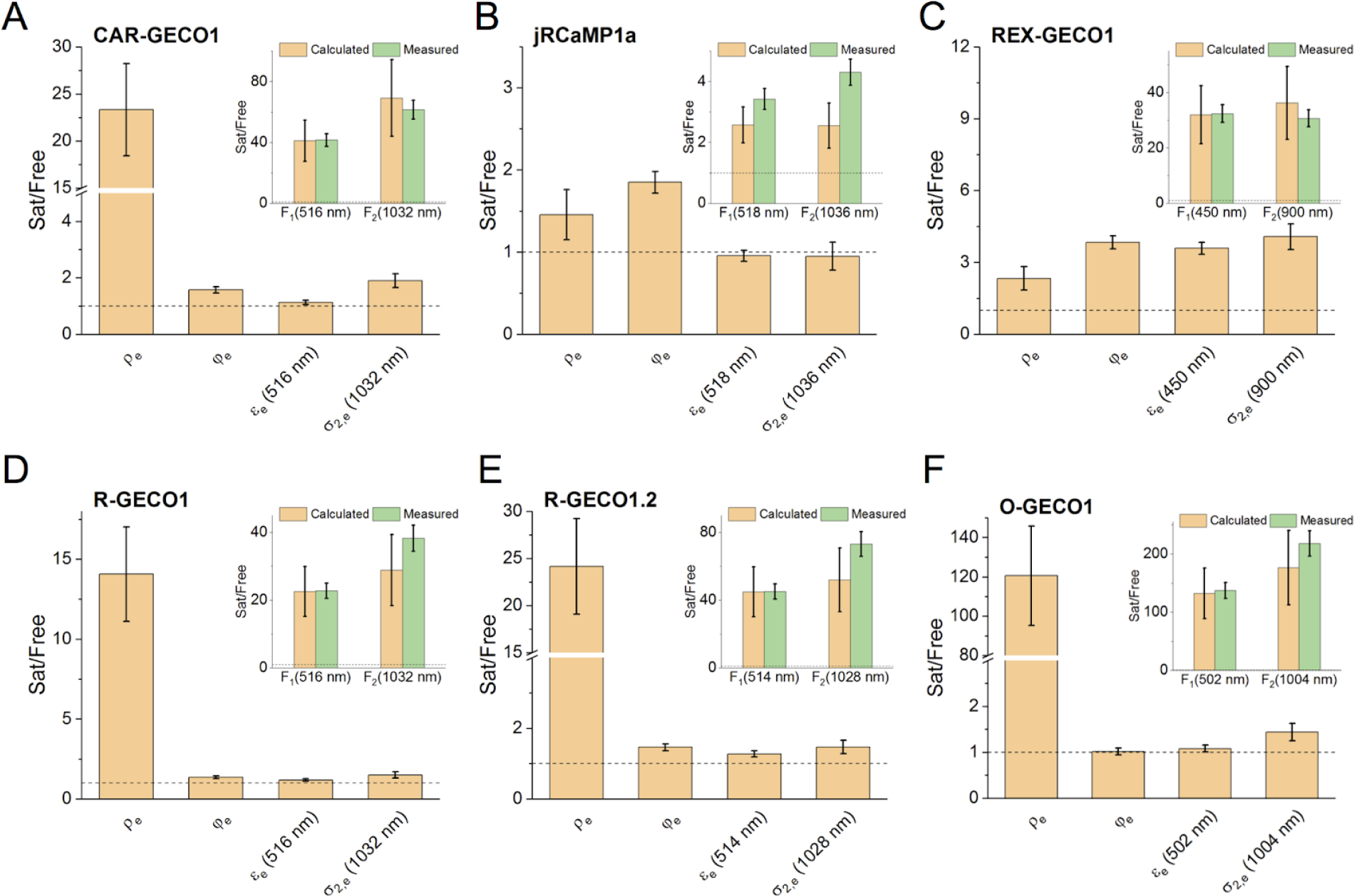
Ca^2+^-saturated/Ca^2+^-free (Sat/Free) ratios for ϱ_e_, ϕ_e_, ε_e_, and σ_2,e_ at the excitation wavelength indicated by ϱ_e_ ϕ_e_ the vertical dashed lines in Figures S4 and S5. Insets: Overall Ca^2+^-saturated/free F1 and F2 ratios, calculated from the independent measurements of ϱ_e_, ϕ_e_, ε_e_, and σ_2,e_ (tan) and directly measured (green). The horizontal dashed line marks a Sat/Free ratio of 1 (no change) for each plot.

**Figure S7.**
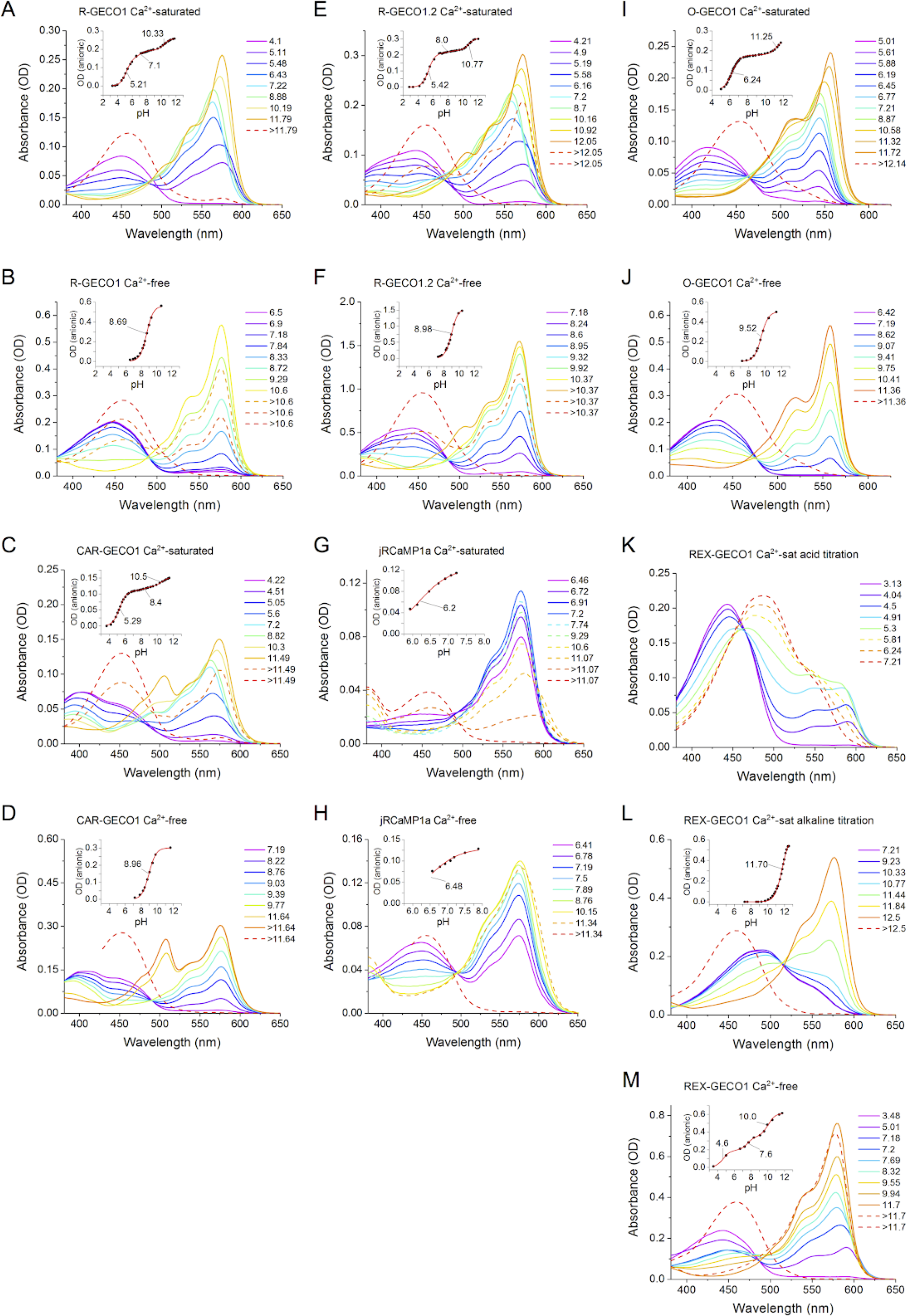
Absorbance pH titrations and p*K_a_*s (*insets*) for R-GECO1, R-GECO1.2, O-GECO1, CAR-GECO1, jRCaMP1a, and REX-GECO1 measured in either Ca^2+^-free or Ca^2+^-saturated buffer. Each spectrum was measured at the pH value designated in the legends. The solid spectra follow the increase of the anionic form peak, while the dashed spectra follow the decrease. Except for (*K*), the final spectrum (red dashed line) belongs to the denatured chromophore. (*L, M*) For clarity, Ca^2+^-saturated REX-GECO1 pH titration are split into the acid and alkaline titrations, respectively. (*Insets*) Fitted p*K_a_* values are indicated on the curve. Note that for each protein, the Ca^2+^-saturated and Ca^2+^-free plots have an equal pH scale.

